# Nephron segmentation and patterning in kidney organoids can be modulated by distinct FGF subfamily members

**DOI:** 10.1101/2025.06.10.658766

**Authors:** Allara K. Zylberberg, Emma I. Scully, Pei Xuan Er, Hannah Baric, Michelle Scurr, Mian Xie, Thanushi Peiris, Sara E. Howden, Kynan T. Lawlor, Melissa H. Little

## Abstract

Human pluripotent stem cell-derived kidney organoids replicate embryonic nephrogenesis in a 3D-culture system. Recent advances suggest that refining the culture environment to replicate spatiotemporal cues present during embryonic organogenesis improves patterning. Here, this paradigm was applied to FGF signalling, a key regulator of embryonic nephron progenitor maintenance, nephrogenesis and ureteric branching. Both FGF8b and FGF10 signalling is sufficient to support nephrogenesis, with each having distinct effects on nephron patterning. FGF10 enhanced the initial WT1+ mesenchymal population, leading to proximally biased nephrons, while FGF8b biased toward early distal patterning, leading to the formation of cells with connecting segment identity. The addition of both FGF8b and FGF10 together had an additive effect, leading to a balance of proximal and distal patterning. This differential patterning was retained in tissue transplanted under the murine renal capsule, with FGF8b-treated organoids displaying increased distal/connecting segments. These findings highlight plasticity during organoid nephrogenesis that can be modulated by FGF signalling and identify an approach to refine nephrogenesis toward key cell types.

## Introduction

Nephron function depends on morphologically and functionally specialised segments that are precisely positioned along the proximo-distal nephron axis. Each nephron originates from mesenchymal nephron progenitors, which form an epithelial precursor structure that undergoes growth and segmentation to form spatially restricted, specialised cell types (1). Human kidney organoids are generated from pluripotent stem cells by mimicking the embryonic specification of intermediate mesoderm, giving rise to nephron progenitors. When aggregated in 3D culture, these progenitors form nephrons with proximo-distal patterning and specialised cell types that mirror embryonic nephrogenesis (2–4).

In the embryo, nephron progenitors form a niche with the neighbouring ureteric tips, providing reciprocal signals that maintain progenitor self-renewal, drive tip growth and initiate nephrogenesis (1). As new nephrons grow and elongate, proximo-distal patterning is established though a complex cascade of signalling events, with mouse studies indicating key roles for Notch, Wnt and BMP signalling *in vivo* (5, 6). While key signalling events in nephron initiation and patterning appear to be recapitulated in stem cell-derived organoids, these nephrons form in the absence of a niche, relying on self-organisation driven by minimal external cues (3, 7). Recent studies indicate that incorporating key aspects of the embryonic nephrogenic signalling can improve or modulate organoid models (4, 8).

The FGF superfamily is comprised of 22 ligands within 6 subfamilies that signal through four receptors (FGFR1, 2, 3 and 4) and a truncated receptor (FGFR1L) (9). FGF ligands and receptors have been reported as critical for the formation of the early metanephric kidney, including ureteric bud formation/branching and the support of nephron progenitor populations during nephrogenesis (10–13). During kidney development, FGF8 is first expressed just after nephron formation in the distal renal vesicle. Here, FGF8 is downstream of WNT4 expression with loss of either of these genes negatively impacting completion of nephrogenesis (14, 15). The specific expression in the distal nephron might suggest a differential role in distal segment identity and connection of the nephron to the adjacent ureteric tip. At an earlier point in embryonic development, FGF8 expression in the presomitic mesoderm promotes axial extension of the nephric duct, ensuring progression of urinary tract development (16, 17). Hence, this growth factor may encourage cellular migration, a feature of the distal renal vesicle during invasion of the adjacent ureteric epithelium (18). FGF9 is expressed in the nephron progenitors of the cap mesenchyme, where it is required for nephron progenitor self-renewal (19). Knockout of FGF9, in conjunction with knockout of another subfamily member, FGF20, results in renal agenesis (19).

The metanephric mesenchyme secretes FGF10, a member of the FGF7 subfamily, which binds to FGFR2b in the ureteric bud to induce branching. Knockout of FGF10 results in fewer ureteric tips and hence smaller kidneys (20). Conversely, overexpression of FGF10 can compensate for the loss of GDNF by maintaining branching in the absence of GDNF-RET signalling (21). Indeed, FGF10 appears to signal through ETV4/5 as does GDNF, the primary regulator of ureteric branching (21). In zebrafish, the initiation of neonephrogenesis involves FGF10 and FGF4 which serve to induce the formation of aggregates from which these new nephrons arise (22). A number of other FGF superfamily ligands are expressed in the cap mesenchyme (FGF1, 7, 9, 10 and 20), ureteric epithelium (FGF9), renal vesicle (FGF8) and ureteric bud (FGF12) (9).

Stem cell-derived models of the human kidney (kidney organoids) show the formation of early nephrons from a nephron progenitor population (2, 3). This study examined whether modulating FGF signalling, one of the key developmental pathways involved in kidney development, could lead to improvements in a stem cell-derived organoid system. Using a combination of transcriptional profiling, reporter stem cell lines and 3D imaging across time, distinct shifts in nephron segmentation were achieved that remained stable after organoid transplantation. FGF8b promoted distalisation of forming nephrons, whereas FGF10 promoted WT1 expression in early nephron and ultimately improved patterning to glomeruli. The expansion of both proximal and distal nephron segments, including the formation of a connecting segment, was enhanced via the combined presence of FGF10 and FGF8b during the initiation of nephron formation, with this shift in patterning shown to be stable after organoid transplantation.

## Results

### Differential nephron formation and patterning shifts in response to FGF ligands

The kidney organoid protocol deployed in this study was based on the previously described kidney micro-organoids in which aggregation and nephron patterning occurs in suspension (2). Kidney organoids were generated using a 7-day monolayer culture to pattern to intermediate mesoderm and a further one day after organoid aggregation (D7+1), at which point growth factors were switched from FGF9 into either FGF10, FGF8b or a combination of both growth factors (FGF10+FGF8b 50:50) (Figure 1A). Kidneys were assessed early in organoid development (renal vesicle stage; D7+5) and later stages (D7+11 and D7+18) after patterning and segmentation had progressed (Figure 1B). A previously characterised double reporter line, MAFB^mTagBFP2^;GATA3^mCherry^ human iPSC line (23) was used to generate kidney suspension organoids, allowing the visualisation of MAFB (podocytes) and GATA3 (distal nephron epithelia).

**Figure 1:**
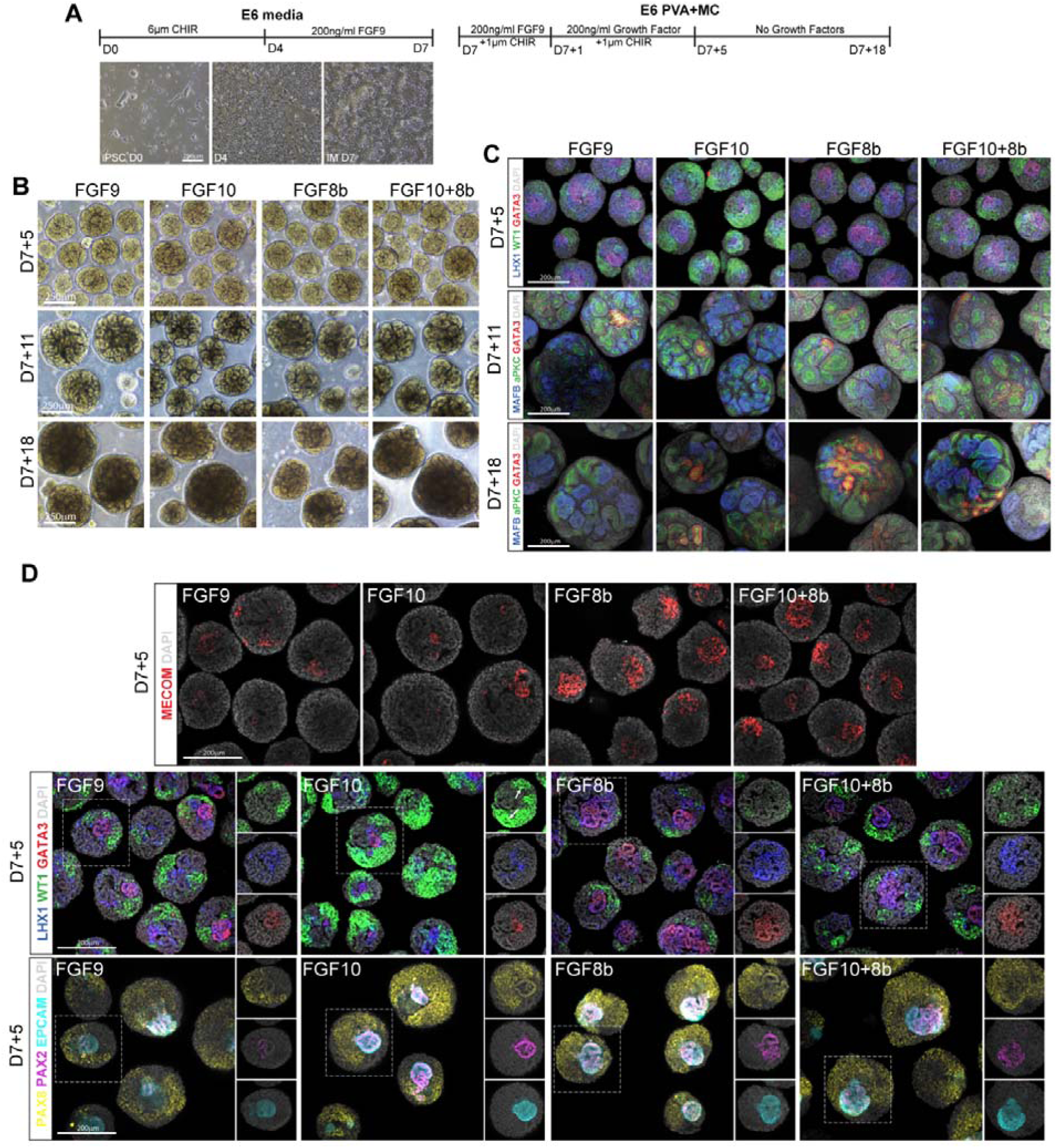
Modifying growth factor conditions alters nephron patterning in suspension kidney organoids. **A.** Schematic depicting differentiation protocol, where Growth Factor refers to either FGF9, FGF10, FGF8b or FGF10+8b (100ng/ml), including representative brightfield images of monolayer differentiations. Scale bar = 250 µm. **B.** Representative brightfield images of organoids following variations at multiple timepoints. **C.** Representative maximum projection confocal immunofluorescence images of kidney organoids using the human iPSC line MAFB^mTagBFP2^;GATA3^mCherry^. At D7+5, immunofluorescence depicts early distal nephrons (GATA3; red), renal vesicle (LHX1; blue) and early proximal nephron (WT1; green) and a nuclear marker (DAPI; grey). At D7+11 and +18, it shows podocytes in the glomeruli (MAFB; blue), late distal tubule (GATA3; red), apical surface of the renal tubules (aPKC; green) and a nuclear marker (DAPI; grey). Scale bars = 200 µm. **D.** Confocal immunofluorescence images of kidney organoids at D7+5 showing an early distal marker (MECOM; red), early distal nephrons (GATA3); early proximal nephron (WT1; green) and early kidney markers (PAX2; magenta, PAX8; yellow), an epithelial marker (EPCAM; cyan) and a nuclear marker (DAPI; grey). Arrows represent early proximal marker WT1 in the developing renal tubules as opposed to the surrounding mesenchyme. Scale bars = 200 µm.

Organoids grown in all conditions displayed appropriate renal patterning and segmentation, containing glomeruli (including podocytes) and distal epithelium as assessed via immunostaining (Figure 1C, D). All organoids displayed appropriate early renal markers regardless of FGF ligand early in development (PAX8, PAX2), and these were beginning to epithelialise (EPCAM) as expected (Figure 1D). At D7+5, an increase in mesenchymal WT1 expression was seen with FGF10 (Figure 1C, D), suggesting enhanced progenitor patterning, although a proportion of WT1-positive cells also appeared to have commenced nephron formation (Figure 1D, white arrows in FGF10). Although markers at D7+5 showed no differences of the late distal marker GATA3, analysis of a marker of distal tubules (MECOM) showed differential patterning, with FGF8b and FGF10+8b having increased expression (Figure 1D).

Automated imaging capture of large fields of organoids enabled quantification of proximal (WT1 early, MAFB late) and distal (GATA3) nephron segments per organoid (Supplementary Figure 1A). WT1 percentage per organoid was increased in FGF10 at D7+5 (Figure 2A), and an increase in GATA3 percentage per organoid was evident in organoids cultured with FGF8b was seen at both D7+11 and +18 (Figure 2A). This was also true for the FGF10+FGF8b condition at D7+18, presumably in response to the presence of FGF8b (Figure 2A). Proximal patterning, as assessed by MAFB staining, was slightly reduced in the FGF10+FGF8b condition compared to FGF9, however was less evident by D7+18 suggesting a slight delay in glomeruli formation in the presence of FGF8b (Figure 2A). Overall, this indicates that FGF8b increased the proportion of distal segments in kidney organoids, whereas FGF10 formed a more developed cap mesenchyme-like population with more proximal nephrons early in renal development. All observations were validated in biological replicate experiments and these findings were consistent when the same assessment was performed in an independent human iPSC human line (Supplementary Figure 2).

**Figure 2:**
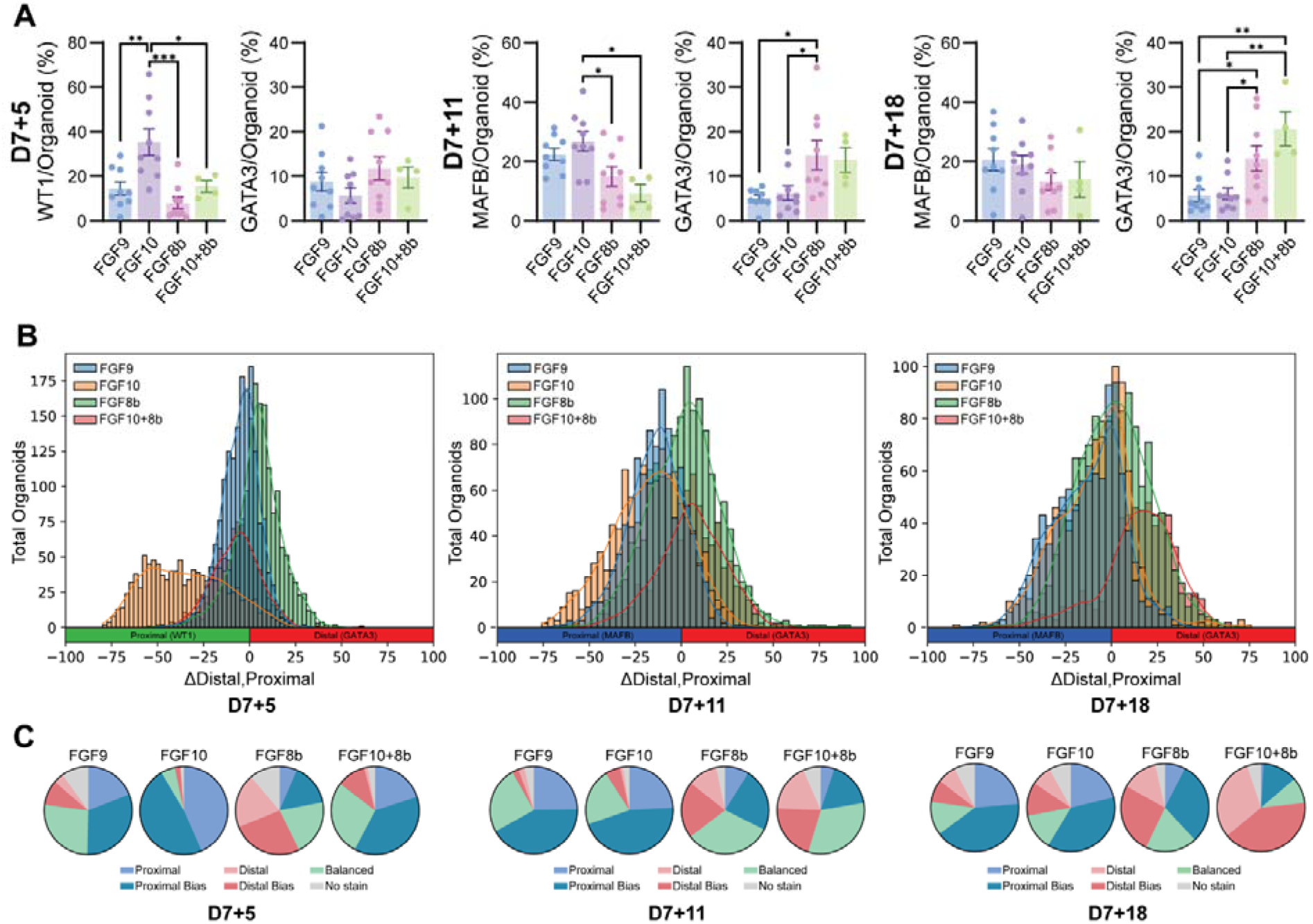
Kidney organoids in suspension show heterogeneity in patterning per organoid with shifts in patterning reflecting a change in the percentage of organoids with a distinct morphology. **A.** Quantification of the proportion of positive segments identified by immunofluorescence in the average total organoid area per organoid. One replicate is the average percentage of at least 45 individual organoids per differentiation. Analysis was undertaken at multiple timepoints. **B.** Histograms showing the distribution of pooled organoids from each differentiation per condition, where organoids to the left of 0 have a proximal bias and the right shows a distal bias. **C.** Pie charts showing pooled organoids subclassified based on morphology. All graph data indicate mean ± SEM; *n=8-9* individual differentiations per condition per time point, except for FGF10+8b where *n=4*. A one-way ANOVA was used for all analyses, using a Tukey’s multiple comparisons *post hoc* test (D) (\**P*□≤□0.05, \*\**P*□<□0.01).

### FGF ligands impact patterning heterogeneity between organoids

The nature of these experiments is such that each organoid is independently undergoing morphogenesis. This has previously been proposed to vary depending upon the diameter of the individual aggregates, with a previous protocol sieving to remove non-patterning suspension culture micro-organoids less than 200µm or greater than 500µm (24). In order to evaluate whether variations in growth factor genuinely impacted patterning or altered micro-organoid heterogeneity, the average size and patterning morphology of organoids in all batches across time were evaluated. An examination of kidney organoid size showed no change compared to FGF9 at any time point (Supplementary Figure 1D).

To quantitatively assess the impact on patterning, organoids were analysed within batches to identify the percentage of distal (GATA3) and proximal (MAFB) segments per organoid across a minimum of 45 organoids assessed per differentiation, per growth factor (Supplementary Figure 1A). To assess overall shifts in patterning, the difference of total distal area (as identified by percentage GATA3) and proximal area (as identified by either percentage WT1 at D7+5 or percentage MAFB at D7+11/18) per organoid was calculated, where a difference of zero indicates balanced morphology (Figure 2B). This simple metric allowed pooling of organoids across all batches so shifts in patterning could be identified per condition. At D7+5, there was a strong proximal population when organoids were grown in FGF10, however this stabilised at later timepoints by which time the ligand is not present. At D7+11, there was a distal bias in both FGF8b and FGF10+8b combined, whereas FGF9 has a proximal bias. By D7+18, organoids appeared to have balanced morphology, except for FGF10+8b which had a distal bias (Figure 2B).

To validate the shifts in patterning, organoids were classified based on morphology into one of 5 categories; proximal, proximal bias, balanced, distal bias or distal (Figure 2C; Supplemental Figure 1C). Using the approach, at D7+5, FGF9, FGF8b and FGF10+8b showed balanced morphology, although a distal bias with FGF8b and a proximal bias with FGF9 were now evident. FGF10 had the strongest proximal shift, lacking both balanced and distal phenotypes. By D7+11, FGF9 and FGF10 maintained their proximal bias with minimal distal biases, while FGF8b and FGF10+8b showed increased balance with a similar proportion of distal and proximal bias phenotypes. By D7+18, FGF9 and FGF10 maintained a proximal bias, FGF8b was balanced and maintained an even distal/proximal bias, whereas FGF10+8b shifted toward a distal bias with fewer proximal organoids (Figure 2C, Supplementary Figure 2B).

### Differential proximodistal patterning is evident at initial nephron formation

To examine whether morphological shifts in patterning correspond to changes in cell identity at the transcriptional level, single cell RNA-sequencing was performed on kidney micro-organoids using the MAFB^mTagBFP2^;GATA3^mCherry^ human iPSC line (23) (Supplementary Figure 3A, 5A). To reduce variability, libraries were created from the same monolayer thawed on the same day and analysed at two time points (D7+5 and D7+11).

Clustering of single cell transcriptomes at the nephron formation timepoint (D7+5) identified 14 clusters (Supplementary Figure 3C). Cluster identity was assigned via manual identification of canonical marker genes (Supplementary Figure 3D) and based on a pre-trained classification tool, *DevKidCC* (25). Final curated identities contained several subclusters of metanephric mesenchyme (MM), early nephron (EN) and two clusters identified as early distal tubule (ED1 and ED2) (Supplementary Figure 3C). All clusters were represented regardless of growth factor (Figure 3A), and relative proportion of clusters between growth factor conditions varied. FGF10 induced a higher proportion of early nephron, but the lowest proportion of both ED1 and ED2 (Figure 2B, Supplementary Figure 2E). FGF8b showed more ED1 than ED2 (Figure 2B, Supplementary Figure 3E), with this cluster showing higher *EPCAM* expression (Supplementary Figure 4A) and less closely aligned to early nephron than ED2.

**Figure 3:**
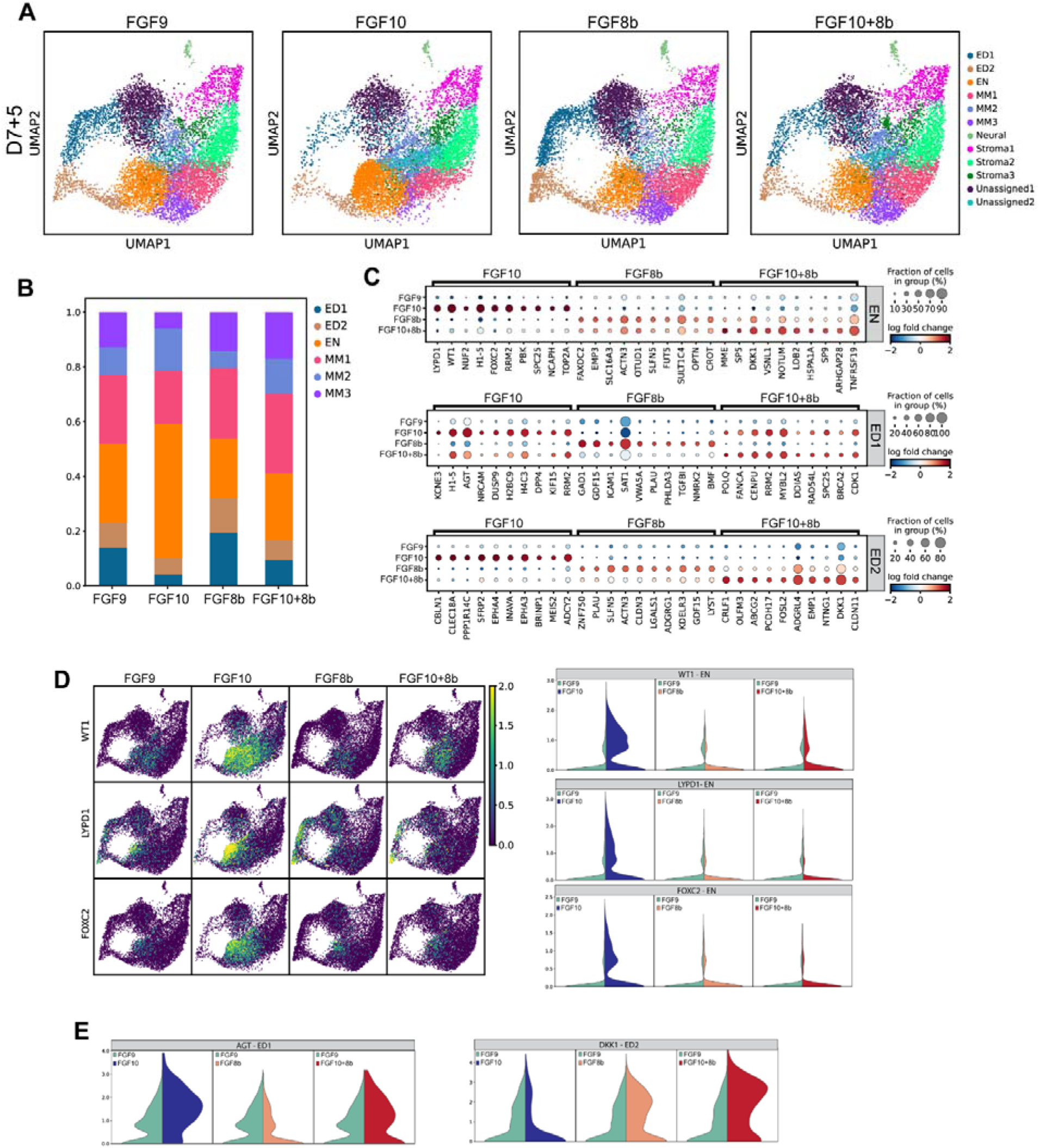
Transcriptional variations in response to FGF ligand in early kidney organoids. **A.** Suspension culture kidney organoids generated from human iPSCs grown in different growth factors (FGF9 control, FGF10, FGF8 and FGF10+8) processed for single cell analysis. UMAP plots of D7+5 organoids identified 14 Leiden clusters. Clusters were identified using marker genes. Population abbreviations: metanephric mesenchyme (MM), early distal (ED), early nephron (EN). **B.** Stacked bar plots showing the proportion of cell types in kidney organoids (nephron specific only). Population abbreviations as above. **C.** Dot plots showing an unbiased analysis of the top 10 significantly differentially expressed genes, expressed with a cutoff of >25% cells having positive expression. These are separated out into cell type clusters, showing early nephron (EN), early distal cluster 1 (ED1) and 2 (ED2). **D.** UMAPs showing profiles of early nephron markers (WT1, LYPD1, FOXC2), which are also depicted as violin plots showing expression of genes identified compared to FGF9. **E.** Violin plots denoting unbiased upregulated genes identified in **(C)**.

Unbiased analysis of the top 10 most differentially expressed genes across all clusters identified an increased expression of WT1 in all metanephric mesenchyme populations when grown in FGF10, in line with image-based patterning analysis (Figure 2, Supplementary Figure 3F). There was also upregulation of *WT1, LYPD1* and *FOXC2* in the early nephron population (Figure 3C, D). An examination of known markers involved in kidney development identified an increase in the expression of markers of the metanephric mesenchyme (*EYA1, SALL1, NR2F2, OSR1*) and nephron progenitors (*SIX1, SIX2, LYPD1, WT1*) (Supplementary Figure 4A). The EN population in response to FGF10 also showed increased expression of pretubular aggregate/renal vesicle (PTA/RV) markers (*WNT4, LHX1*) in the FGF10 early nephron population (Supplementary Figure 4B). An early marker for podocyte development (*FOXC2*) is also upregulated. Both early distal populations have low expression of medial and distal early nephron markers, however markers of the early proximal nephron are upregulated in both of these populations (Supplementary Figure 4B). Although the smallest population compared to the other FGFs, changing from FGF9 into FGF10 appears to result in a higher proportion of expressed nephron progenitor cells as well as a shift to a more proximalised renal vesicle population early in organoid development.

Micro-organoids differentiated in the presence of FGF8b displayed the largest proportion of early distal nephron, particularly ED1 (Figure 3A, B). The MM clusters showed increased expression of nephron progenitors in MM3 (*SIX2, CITED1*) (Supplementary Figure 4A). Both ED1 and ED2 showed upregulation of EPCAM in the presence of FGF8b suggesting a more advanced level of epithelialisation (Supplementary Figure 4B). These clusters also showed an increase in distal early nephron markers *POU3F3, MECOM, GATA3* and *SOX9* (Supplementary Figure 4C). Hence, FGF8b at D7+5 induces a more distal nephron compared to FGF9. These findings were consistent with the image-based patterning analysis (Figure 1C, D, Figure 2). There is also upregulation of *TFAP2A* (ED2) and *TFAP2B* (ED1) (Supplementary Figure 4D), both of which have been previously shown to modulate nephron precursor differentiation into distal nephron in the mammalian kidney (26). This trend was also true for the FGF10+8b mix.

Micro-organoids cultured with FGF10+8b displayed a transcriptional signature that combined features of the FGF10 and FGF8b induced changes. The metanephric mesenchyme clusters were similar to FGF10 (Supplementary Figure 4A), however EN gene expression was more like FGF8b, with upregulation of medial *IRX3* and distal (*SOX9* and *DKK1*) nephron markers (Supplementary Figure 4B). This indicates that the early nephron cluster grown in FGF10+8b has a more balanced proximal/distal early nephron population. The proportions of early distal clusters with FGF10+8b is less than FGF9, however expression levels of early distal nephron markers are significantly upregulated in both distal clusters (*POU3F3, MECOM, GATA3, DKK1*) (Supplementary Figure 4C). This indicates that FGF10+8b maintains a proximal early nephron population while also promoting the formation of an early distal population. Unbiased analysis of the top 10 differentially expressed genes identified *CRLF1* in ED2 (Figure 3C, Supplementary Figure 3G). *CRLF1* has been previously shown to be expressed in the ureteric bud tips and acts downstream of GDNF (27). There is also expression of *WNT9b* (Supplementary Figure 3G) in the same region of the UMAP, potentially indicating FGF8b and the FGF10+8b have a small ureteric bud-like population, possibly driven by FGF8. All differentially expressed genes at this timepoint can be found in Supplementary File 1.

### Proximal bias evident in response to FGF10 with distal bias evident from FGF8b and FGF10+8b

To investigate the impact on patterning beyond the early nephron, single cell RNA-sequencing was also performed on all conditions at D7+11 (organoids shown in Supplementary Figure 5A). All clusters were represented under all conditions with minimal shifts in the proportion of each cluster between conditions (Figure 4A). Based on imaging, all conditions displayed normal renal patterning, albeit FGF9 and 10 had a proximal shift, including the development of a strong proximal tubule population with FGF10 (LTL+), whereas FGF8b and FGF10+8b had a more balanced morphology (NPHS1+ and ECAD+) (Figure 4F).

**Figure 4:**
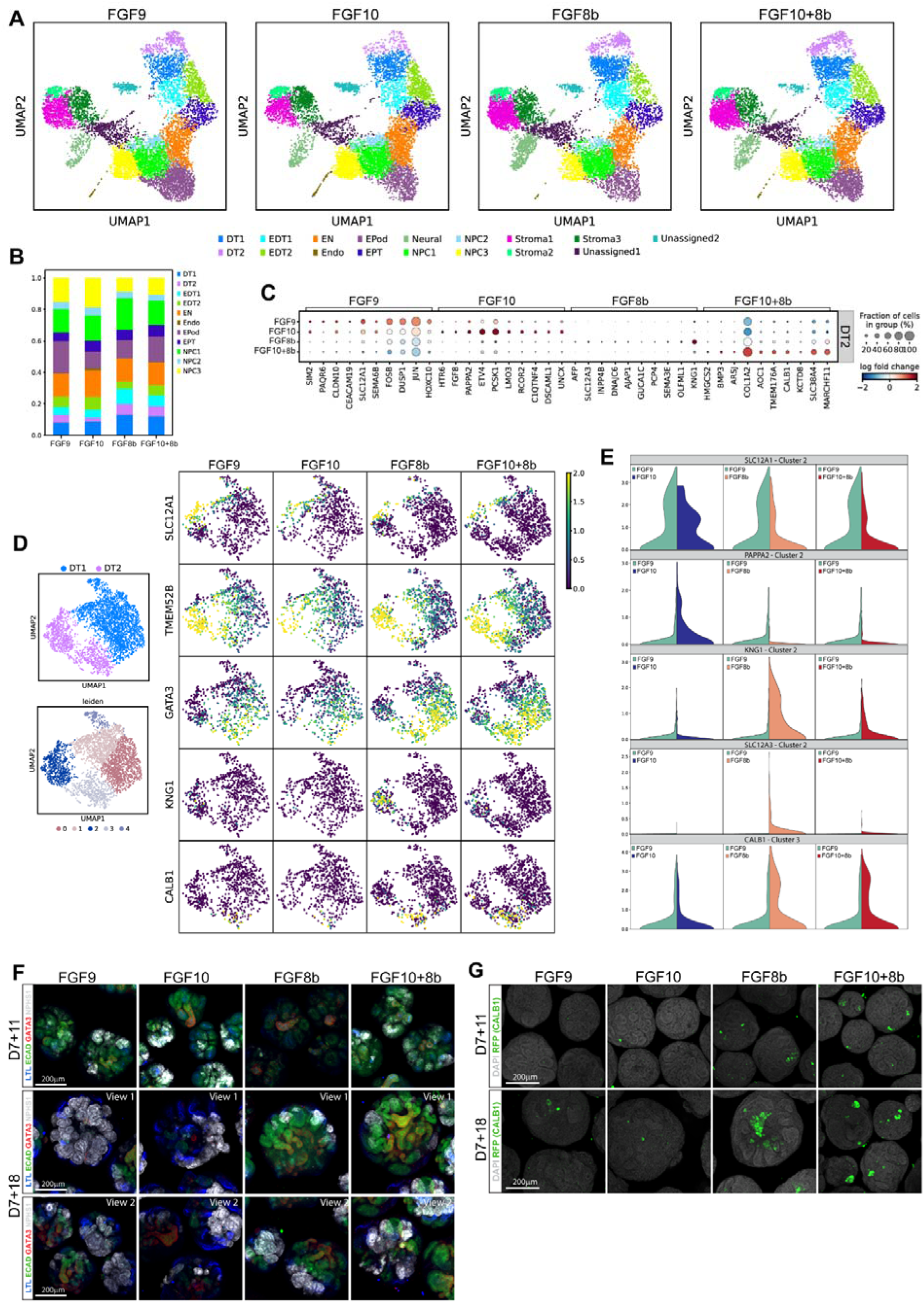
Single cell characterisation identifies the presence of a connecting segment in late kidney organoids. **A.** Suspension culture kidney organoids generated from human iPSCs grown in different growth factors (FGF9 control, FGF10, FGF8 and FGF10+8) processed for single cell analysis. UMAP plots of D7+11 organoids identified 17 Leiden clusters. Clusters were identified using marker genes. Population abbreviations: distal tubule (DT) separated into 2 Leiden clusters (DT1, DT2), early distal tubule separated into 2 Leiden clusters (EDT1, EDT2), early nephron (EN), endo (endothelial), early podocytes (EPod), early proximal tubules (EPT), nephron progenitor cells (NPC) separated into 3 Leiden clusters (NPC1, NPC2, NPC3), Stroma separated into 3 Leiden clusters and a neural population. **B.** Stacked bar plots showing the proportion of cell types in kidney organoids (nephron specific only). Population abbreviations as above. **C.** Dot plots showing an unbiased analysis of the top 10 significantly differentially expressed genes per growth factor in distal tubule cluster 2 (DT2), separated by growth factor. **D.** Both distal clusters separated re-clustered into 5 Leiden clusters. Based on UMAPs, these have biases towards different segments of the distal nephron, including a connecting segment. **E.** Violin plots showing expression of genes in varied growth factors (FGF10, FGF8b, FGF10+8b) compared to FGF9. These are Loop of Henle (SLC12A1), macula densa (PAPPA2), distal convoluted tubule/connecting segment (SLC12A3, KNG1) and connecting segment (CALB1). **F.** Represented Immunofluorescence (maximum projection) of D7+11 and +18 organoids grown in different growth factors. Images depict proximal tubules (LTL; blue), distal tubules (ECAD, green), late distal tubules (GATA3; red) and podocytes (NPHS1; grey). Scale bar = 200 µm. **G.** Immunofluorescence (maximum projection) of CALB1-mCherry reporter derived D7+11 and D7+18 organoids grown in different growth factors. Images depict connecting segment (CALB1; green). Scale bar = 200 µm.

When data was subsetted and re-clustered to examine renal cell types only, the relative proportion of clusters between growth factor conditions did vary slightly (Figure 4B), with FGF9 having the highest proportion of early podocytes (20.3%), whereas FGF8b had the highest proportion of distal tubules (DT1+DT2 – 20%) (Figure 4B). When examining gene expression changes, DT2 had the most variation, and was thus the focus for subsequent analysis of the D7+11 data.

### FGF8 and FGF10+8b pattern to connecting segment and distal convoluted tubule

Unbiased analysis comparing the top 10 differentially expressed genes between conditions in DT2 showed that changing growth factors reduced the proportion of cells with a thin descending LOH-like identity that formed with FGF9 (*SIM2, SLC12A1*). Of note, FGF10 upregulated macula densa genes (*ETV4, PAPPA2*) in this population, as did FGF10+FGF8b (*TMEM176A, BMP3*). FGF8b upregulated genes associated with the distal straight tubule (KNG1) as well as distal convoluted tubule (*SLC12A3*), and distal tubule (*OLFML1*) (Figure 4C). All three of those genes are also present in the connecting segment, and there is also upregulation of CALB1 (strongly expressed in the nephron connecting segment). FGF10+8b cells also had increased expression of connecting segment markers (*CALB1, KCTD8, ARSJ*) (Figure 4C). Differentially expressed genes can be found in Supplementary File 2.

A re-clustering of the distal tubule clusters, DT1 and DT2, identified 5 subclusters with broad expression of distal tubule/ loop of Henle / ureteric epithelial markers. Clusters 2 and 3 were more represented in FGF8b and FGF10+FGF8b organoids, with cluster 2 showing expression of genes previously associated with distal convoluted tubule markers and connecting tubule (*SLC12A3, KNG1)* while cluster 3 expressed the connecting segment gene *CALB1* and SCNN1G. Clusters 0, 1 and 2 all showed distal tubule markers (*MECOM, MAL, GATA3, TMEM52B*), with cluster 2 most like the thin descending Loop of Henle (*SLC12A1, MUC1, SPP1, KCNJ1*) (Supplementary File 3). Taken together, the unbiased analysis and re-clustering indicates that micro-organoids cultured with FGF8b and FGF10+8b had a smaller loop of Henle population, but in exchange created a distinct CALB1+ connecting segment. FGF10 and FGF10+8b developed a macula densa population (Figure 4C, D, E). As a result, FGF10 likely promotes formation of a macula densa, whereas FGF8b promotes formation of the connecting segment. The presence of a connecting segment population was confirmed via immunofluorescence (Figure 4G) using a CALB1-mCherry reporter line. A full list of genes can be found in Supplementary File 3.

### Proximo-distal patterning bias is maintained *in vivo*

To identify if variations in growth factors maintain their morphology long term, D7+5 MAFB^mTagBFP2^GATA3^mCherry^ reporter line-derived organoids were transplanted in conjunction with their corresponding growth factor under the renal capsule of female immunocompromised NSG mice (28). After 14 days, whole host kidneys with associated graft were embedded in 3% low melting point agarose and sliced into 600µm vibratome sections and analysed using 3D imaging with key patterning markers to mark glomeruli (NPHS1) and distal tubules (GATA3, using a GATA3^mCherry^ reporter) (Figure 5A). Organoids were transplanted at D7+5 as these are early in nephron development and show the most transcriptional differences with modified growth factors. These organoids are also less committed to specific tubule types.

**Figure 5:**
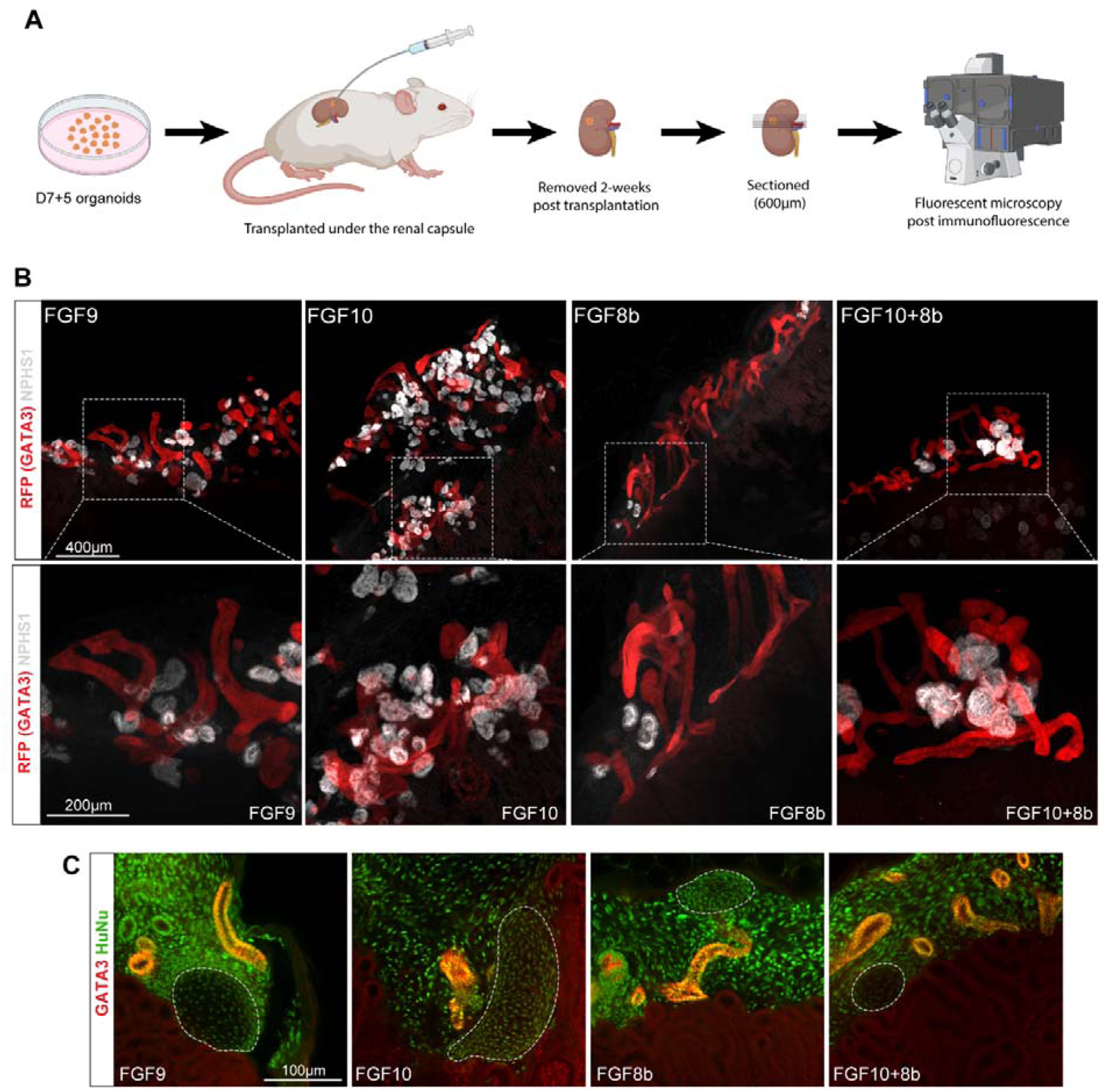
Differential patterning is maintained post transplantation. **A.** Schematic representation of transplant experimental design. Organoids at D7+5 grown in various growth factor conditions (FGF9, FGF10, FGF8b, FGF10+8b) were transplanted under the renal capsule of mice for 2 weeks, at which point animals were ethically euthanised and samples were sectioned in 600um sections, underwent immunofluorescence and were imaged on a spinning disk confocal microscope. **B.** Representative images of graft tissue patterning, depicting a distal tubule marker (RFP - GATA3 in human derived tissue only; red) and glomeruli (NPSH1; grey). Scale bar = 400µm. Higher magnification images of (b), showing distal tubules (RFP – GATA3; red) and glomeruli (NPSH1; grey). Scale bar = 200µm. **C.** Off-target populations in kidney organoids develop into cartilage, predominantly in the condition grown in the FGF10 ligand. Image depicts human derived tissue (Human nuclear; green) and distal tubules (GATA3; red).

Morphological analysis identified that the early changes in FGF ligand-mediated patterning are maintained long term *in vivo*. Transplantation with FGF9 formed a relatively balanced proportion of distal and glomerular segments. However, while glomeruli formed glomerular tufts, the distal tubules in these population appeared short and disorganised. FGF10 promoted the formation of glomeruli, but the distal tubules maintained a similar phenotype to FGF9, which is unexpected due to the anticipated role for this ligand in branching morphogenesis rather than glomerular patterning (20). Conversely, transplantation of FGF8b promoted the formation of distal tubules that had an elongated phenotype, but these lacked a strong population of glomeruli (Figure 5B). This is likely due to the reduced population of WT1 and other early glomerular markers at D7+5. Transplantation with organoids grown in FGF10+8b had formation of both glomeruli and distal tubules (Figure 5B). Although transplanted organoids developed the expected key renal cell types, there was also sporadic growth of cartilage-like tissue (Figure 5C) which were specifically derived from human organoid tissue. Cartilage developed in all conditions regardless of growth factor, in line with previous subcapsular transplantation of kidney organoids (28).

The glomerulus and distal tubule area was quantified, as was the total glomeruli number. Distal analysis was limited to human organoid derived tubules as immunostaining was based on an antibody specific to the mCherry (GATA3) reporter, that did not stain murine host tubules. As previously noted, there was an increase in total glomeruli number (with an accompanying increase in glomeruli area) in the FGF10 condition. FGF8b had a decrease in glomeruli area, but no significant change in glomeruli number, indicating that glomeruli grown in FGF8b ligand are smaller (Figure 6A, B). The distal tubule area was significantly reduced in FGF10 and FGF8b compared to FGF9, although FGF10+8b displayed no significant difference in glomeruli and distal tubule area. Transplants grown and transplanted in FGF8b or FGF8b/FGF10 developed elongated distal structures that extended towards the host kidney (designated as “touchdowns”), with this likely to be driven by FGF8b (Figure 6B, C). Although 33 separate transplantations of 600µm thick sections were examined, no instances were observed where the graft nephrons showed patent connections with the underlying host kidney, regardless of growth factor condition.

**Figure 6:**
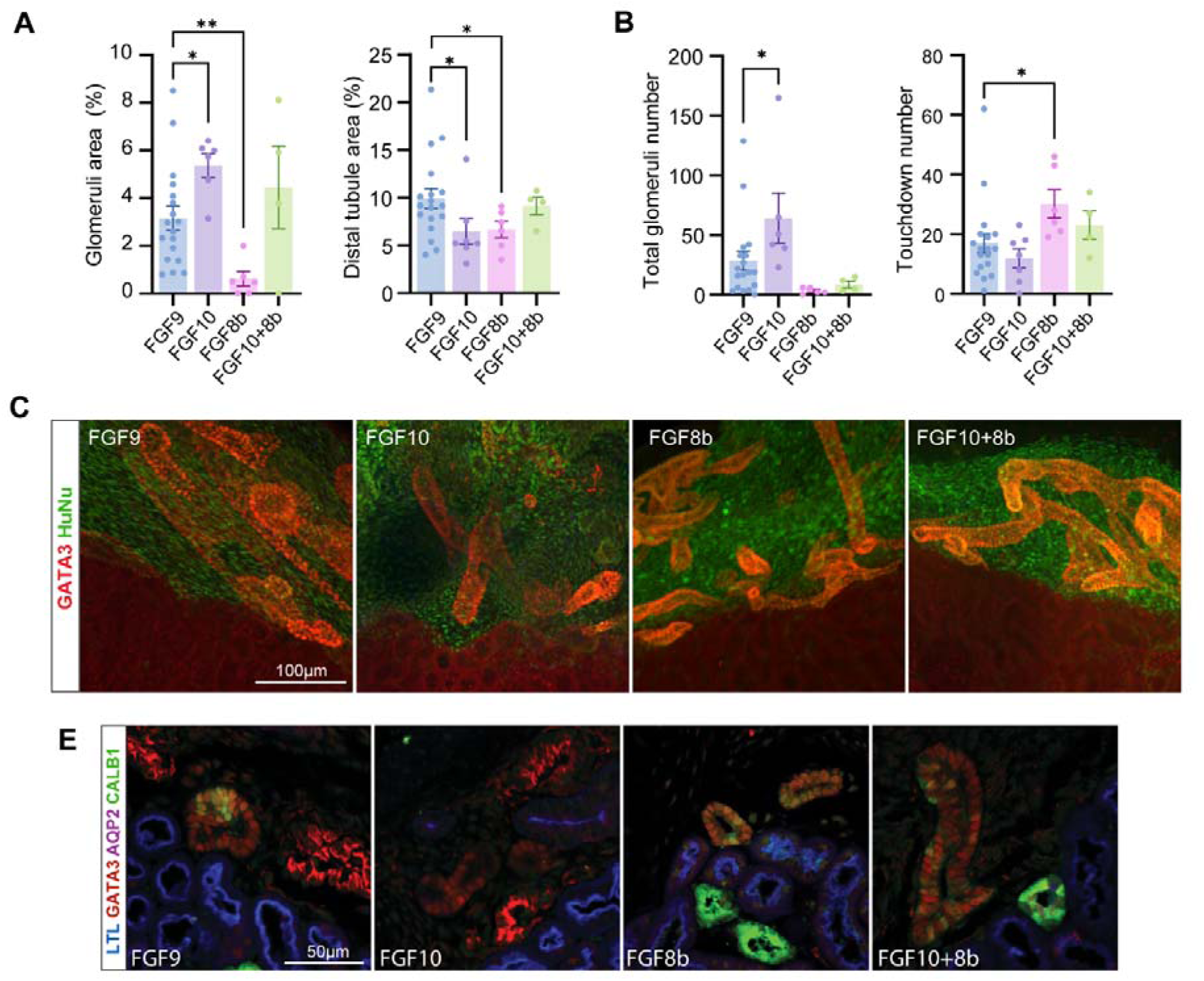
Organoids in FGF8b ligand has increased proportion of organoids touching mouse tubules, and these are CALB1+. **A.** Analysis of total glomeruli and distal tubule area, presented as total percentage of graft area. **B.** Total number of glomeruli per graft and total number of distal tubules in contact with the host tubule. These are designated as “touchdowns”. **C.** Maximum projection representative immunofluorescence showing “touchdowns” per growth factor condition. Image depicts human derived tissue (Human nuclear; green) and distal tubules (GATA3; red). **D.** Immunofluorescence image of 4µm section grafts 2 weeks post transplantation under the renal capsule. These depict proximal tubules (LTL; blue), distal tubules (GATA3; red), connecting segment (human) and collecting ducts (CALB1; green) and collecting duct principal cells (APQ2; magenta). All graph data indicate mean ± SEM. Sample size varies by growth factor (FGF9 *n=18*; FGF10 *n=6*, FGF8 *n=5*; FGF10+FGF8 *n=4*). Kidney organoids from grafts are from two separate differentiations, except for FGF10+FGF8b which is from 1. A one tailed unpaired students t-test is used for analysis, comparing all individual FGFs to FGF9 (\**P*□<0.05, \*\**P*□<□0.01).

To examine whether the connecting segment develops after transplantation, 4µm sections were stained with the distal marker GATA3, as well as the collecting duct marker AQP2 and connecting segment/collecting duct marker CALB1. Sections were also stained with LTL to more readily identify the mouse tubules (Figure 6D). Consistent with previous findings from organoid characterisation (29), no collecting ducts were identified in transplanted kidney organoids (AQP2 negative). In contrast to *in vitro* patterning analyses, engrafted distal tubules derived from tissue grown in either FGF9 or FGF8b, but not FGF10 alone, displayed small regions of connecting segment in all transplants. Thus, FGF-based patterning shifts are sufficient to modulate final cell identity in mature engrafted tissue.

## Discussion

Our ability to control nephron proximo-distal patterning within kidney organoids remains limited. In this study, we report that FGF8b, previously associated with initial nephron patterning and differentially expressed in the distal renal vesicle, can promote nephron distalisation that includes formation of a connecting segment, in kidney organoids. In contrast, the addition of FGF10 promoted more proximalised nephron patterning, including strong glomerular patterning. When combined, FGF10+FGF8b (50:50) retained glomerular patterning and also formed a connecting segment. This suggests additive roles in balanced nephron segmentation for these two FGF ligands.

The addition of FGF8b resulted in the highest expression of SIX2+ CITED1+ cells in early organoid (D7+5) MM populations, implying a more self-renewing nephron progenitor state. This is in line with a recent study which demonstrated that FGF8 prevents BAX/BAK mediated apoptosis in the cap mesenchyme, promoting the nephron progenitor population (30). Normally, this cap mesenchyme undergoes canonical Wnt mediated mesenchymal-to-epithelial transition to upregulate WNT4, initiating the formation of a pretubular aggregate and subsequently triggering FGF8 expression to complete formation of a renal vesicle. These then act to upregulate LHX1 (15). As such, the addition of exogenous FGF8b may force a more immediate epithelialisation event, thus bypassing this initial pretubular aggregate stage. This may also explain the increased levels of tubular EPCAM in early distal tubules of organoids formed with FGF8b. Subsequent culture in FGF8 promoted the formation of a connecting segment and reduced numbers of glomeruli, suggesting an inability to support proximal nephron patterning. FGF10 treatment promoted the expression of a more WT1+ LYPD1+ nephrogenic mesenchyme population with differential expression of LHX1 in the early nephron (D7+5). In zebrafish, FGF10 induces the aggregation of Lhx1+ renal progenitors during injury-induced neonephrogenesis (22), suggesting a similar role here. Alternatively, the upregulation of WT1 may represent a bias towards formation of a more proximal renal vesicle. By D7+11, and even more prominently at D7+18, FGF10 treated organoids became dominantly proximalised with expansive glomeruli and LTL+ proximal tubules. Even if transplanted at D7+5, before this overt proximalisation had occurred, the resulting organoids showed enhanced numbers of enlarged glomeruli at 14 days post-transplant. Combining FGF10 and FGF8b resulted in a phenotype intermediate between that of each growth factor alone. Distinct subclusters showed either more similarity to FGF10 or FGF8b, with the exception of the early metanephric mesenchyme clusters, which appear to display a unique combinatorial gene expression.

A model of progressive nephron progenitor recruitment has recently been proposed as a mechanism that drives nephron formation in murine and human nephrogenesis (31). This model suggests a time dependent process in which nephron progenitors undergo mesenchymal to epithelial transition in sequence from distal to proximal renal vesicle with distinct endpoint fates dictated by the order of recruitment (31). In line with this hypothesis, our findings suggest that FGF10 organoids maintained a higher proportion of WT1+ mesenchymal cells for longer, potentially endowing them with a proximal bias. On the other hand, FGF8b resulted in a more rapid mesenchymal-to-epithelial transition, leading to a distal patterning bias. Interestingly, FGF10+8b forms an intermediate between the two, forming a cap mesenchyme population that is reminiscent of FGF10, but still maintaining the early distal phenotype promoted by FGF8. These early changes in timing during renal nephrogenesis have varying impacts on the development of the distal tubules, resulting in a more developed distal population not observed in kidney organoids previously (2, 3, 32, 33). These included the introduction of a distal straight tubule (FGF10), distal convoluted tubule (FGF8b), a macula densa population (FGF10 and FGF10+FGF8b), and a connecting segment (FGF8b and FGF10+FGF8b). The upregulation of macula densa markers within the D7+11 DT2 cluster was evident in response to FGF10 and hence also in FGF10+8b. This specialised region of the distal tubule remains close to the forming glomerulus. The clear increase in number of glomeruli also seen in response to FGF10 suggests an influence of this population on the distal tubule patterning.

Not all nephrons in the mammalian kidney show the same patterning and segmentation with this varying between the cortical and juxtaglomerular nephrons (34). Cortical nephrons are more abundant and have shorter loops of Henle that only reach the outer medulla whereas juxtaglomerular nephrons have long loops of Henle with prominent descending and ascending thin limbs (35). In rodents and primates (and presumably humans) the glomerular diameter of juxtaglomerular nephrons is larger (34, 36, 37). Hence it has been proposed that there are two distinct nephron phenotypes with potentially distinct roles and sensitivities to injury (34). In this study, *in vitro* and *in vivo* glomerular diameter appeared to be lower in the presence of FGF8b, with the converse in the presence of FGF10. It is therefore possible that the patterning observed in the presence of FGF10 was more similar to a juxtaglomerular patterning whereas those formed in the presence of FGF8b were more cortical in nature. This also fits with the expression of SLC12A3 (distal convoluted tubule). This may suggest a differential role for FGF signaling across time as the juxtaglomerular nephrons arise first and from a younger nephron progenitor population.

The differential nephron patterning in early organoids (D7+5) observed *in vitro* was maintained for a further 2 weeks post transplantation, when co-transplanted with their corresponding growth factor. FGF10 transplants retained a higher number of glomeruli and total glomerular area while FGF8b transplants had a reduction in glomerular area but not total number (resulting in smaller glomeruli) upon transplantation. Both FGF8b and FGF8b:10 patterned organoids retained distal tubular expression of CALB1, but no evidence of a patent connection to the host was observed. An unexpected finding was that the distal tubules present in organoids grown in and transplanted with FGF8b elongated and extended towards the host murine kidney suggesting a capacity to respond either to the surrounding transplant stroma or a trophic guidance from the host. Previous examination in chick embryos identified that the nephric duct extends towards the presomitic mesoderm during axial elongation in response to FGF8 (16). It is possible that the presence of this growth factor at transplantation also retained a more nephric duct tip-like state as has been described in the extending nephric duct in mouse (17). Understanding this better may assist in the *in vitro* control of nephron architecture and alignment.

A clear challenge with the interpretation of stem cell-derived models of the kidney lies in the assignation of cellular identity to clusters, notably at early stages of organoid formation. The presence of two distinct early distal clusters at D7+5 may represent proximal and distal renal vesicle or different stages of maturation from renal vesicle. The presence of a WNT9B+ population in the presence of FGF8b may even suggest that the most distalised structures contain epithelium with similarity to nephric duct. What is clear is that glomerular maturation is better supported by FGF10, CALB1+ distal epithelium by FGF8 and the combination generates a balanced nephron with a CALB1+ connecting segment able to retain identity for 2 weeks after transplantation. These observations will enable the generation of more distalised nephron models both for studies into distal tubulopathies and in tissue engineering efforts.

## Acknowledgements

This research was supported by the Novo Nordisk Foundation Center for Stem Cell Medicine (reNEW) under the Novo Nordisk Foundation grant number NNF21CC0073729. This research was also funded in part by the National Health and Medical Research Council (GNT2011314). For the purposes of open access, the author has applied a CC BY public copyright licence to any Author Accepted Manuscript version arising from this submission. We acknowledge the Murdoch Children’s Research Institute Flow Cytometry and Imaging Core, the Murdoch Children’s Research Institute Genomics Facility for 10x Gene Expression Flex single cell library preparation and sequencing, and the Murdoch Children’s Research Institute Animal Facility. We would also like to acknowledge Fernando Rossello and Michael See from the reNEW Bioinformatics Hub for advice on the bioinformatics analysis.

## Author contributions

A.K.Z., K.T.L. and M.H.L. contributed to experimental design and planning. A.K.Z., P.X.E., H.B. and M.X. performed experiments and data acquisition. M.S. performed transplantation experiments, and H.B. and T.P. analysed the transplant data. A.K.Z. and E.I.S. performed bioinformatics and image analysis. S.E.H. generated the CALB1-mCherry iPSC line. A.K.Z, K.T.L and M.H.L. wrote the manuscript.

## Materials & Methods

### Lead Contact

Requests for further information and resources should be directed to and will be fulfilled by the lead contact, Melissa H. Little.

### Materials Availability

All cell lines used in this study are available upon request.

### Data and Code Availability

Datasets generated during this study are available at Gene Expression Omnibus, under GEO: GSE299104. Code related to analyses in this study is available at: https://github.com/KidneyRegeneration/Zylberberg2025.

### iPSC line culture and maintenance

All iPSC lines were maintained and expanded at 37°C and 5% CO2 in Essential 8 medium (Thermo Fisher Scientific, Waltham, MA) on 6 well plates coated with Matrigel (BioStrategy, Victoria, Australia). Media was changed daily and passaged every 2-3 days via a 3-minute incubation in EDTA in 1xPBS at 75% confluency. Reporter iPSC lines in this study include MAFB^mTagBFP2^;GATA3^mCherry^ (derived from human foreskin fibroblasts [CCD-1112Sk/CRL-2429^TM^, ATCC] and CALB1^mCherry^ (derived from SCT3010 iPSCs - hPSCreg ID# MCRIi032-A) (3, 45).

### Directed differentiation and generation of suspension kidney organoids

#### Monolayer differentiation

Differentiation of iPSC lines and organoid culture was performed as previously published (3). In brief, iPSCs were dissociated to almost single cell suspension through a 3-minute incubation in TrypLE (Thermo Fisher Scientific) and seeded onto a Laminin-521 (BioLamina, Sundbyberg, Sweden) coated (50uL/mL)) 6-well plate when they reached approximately 75% confluency. 24 hours post seeding, the differentiation was commenced through the addition of 2.5ml TeSR-E6 medium (StemCell Technologies) supplemented with 6uM CHIR99021 (R&D Systems) at a seeding density of 120,000 cells/well. This medium was given for a period of 4 and was refreshed every second day. Four days after the differentiation began, cells were exposed to 200ng/ml of FGF9 (R&D Systems) and 1ug/mL Heparin (Sigma Aldrich, St Louise, MO) in 3mL mTeSR-E6 for 3 days, which was refreshed every second day until the time of organoid aggregation at day 7 of differentiation.

#### Organoid generation

To generate organoids, day 7 differentiated cells were washed with 1.5ml 1xPBS, and dissociated using 1.5ml EDTA for 3 mins at 37°C. EDTA was aspirated and cells were resuspended in TeSR-E6 supplemented with 200ng/ml FGF9, 1ug/ml Heparin, 1uM CHIR99021, 0.1% Poly Vinyl Alcohol (PVA), 0.1% Methyl Cellulose (MC), 10uM Rho kinase inhibitor (In Vitro Technologies) and 0.5% Antibiotic-antimycotic (Life Technologies). Resuspended small clumps were transferred to 6cm^2^ low adhesion plates (Greiner Bio-One), where organoids formed spontaneously after placement on an orbital shaker (Ratek) rotating at 60 rpm at 37°C and 5% CO2. After 24 hours, media was replaced with 1ug/ml Heparin, 1uM CHIR99021, 0.1% PVA, 0.1% MC and 0.5% Antibiotic-antimycotic for a further 4 days. In addition, media contained either 200ng/ml FGF9, 200ng/ml FGF10 (R&D Biosystems) or 200ng/ml FGF8b (R&D Biosystems). From D7+5, organoids were refreshed with TeSR-E6 supplemented with 0.1% PVA and 0.1% MC until the desired end point, refreshing every 2- 3 days.

### Generation of CALB1^mCherry^ iPSCs

A targeting vector encoding a T2A-mCherry cassette flanked by 0.7 kb and 0.8 kb homology arms corresponding to sequences upstream and downstream of the *CALB1* stop codon was introduced into SCT3010 iPSCs (hPSCreg ID# MCRIi032-A), along with mRNA encoding Cas9 and a plasmid encoding a sgRNA specific to the 3’ end of the *CALB1* locus using the Neon Transfection Device. Successfully targeted iPSCs (expressing the mCherry reporter) were purified by two rounds of fluorescent activated cell sorting. Targeted integration was confirmed by PCR analysis using primers flanking the 5’ and 3’ recombination junctions.

### Immunofluorescence and confocal microscopy

Organoids were fixed and stained as previously published (2) with minor amendments. In brief, organoids were collected into a 15ml falcon tube and remaining media was aspirated. Paraformaldehyde 16% aqueous solution (Electron Microscopy Services) was diluted to 4% using 1xPBS and added to the organoids for 10 minutes at room temperature, followed by 1 wash in 0.1% Triton-X 100/PBS (Sigma-Aldrich). For staining and imaging, organoids were moved into a 96 well glass bottom plate. Primary antibodies were diluted in 0.1% Triton-X 100/PBS supplemented with 5% donkey serum and are detailed in Table 1. Primary antibodies were detected with Alexa Fluor-conjugated fluorescent secondary antibodies (Life Technologies) at 1:500. Nuclei were detected with DAPI diluted at 1:1000 in PBS given with the secondary antibody diluent. Organoids were in antibody diluents overnight, with 2 rinses and a 3-hour wash between primary and secondary antibody incubations, as well as after the final secondary antibody incubation. Organoids were mounted in 90% glycerol and imaged on the Andor Dragonfly 500 Spinning Disk confocal microscope (Andor Technology, Belfast, United Kingdom). Images were processed using Fiji ImageJ (38).

**Table 1:**
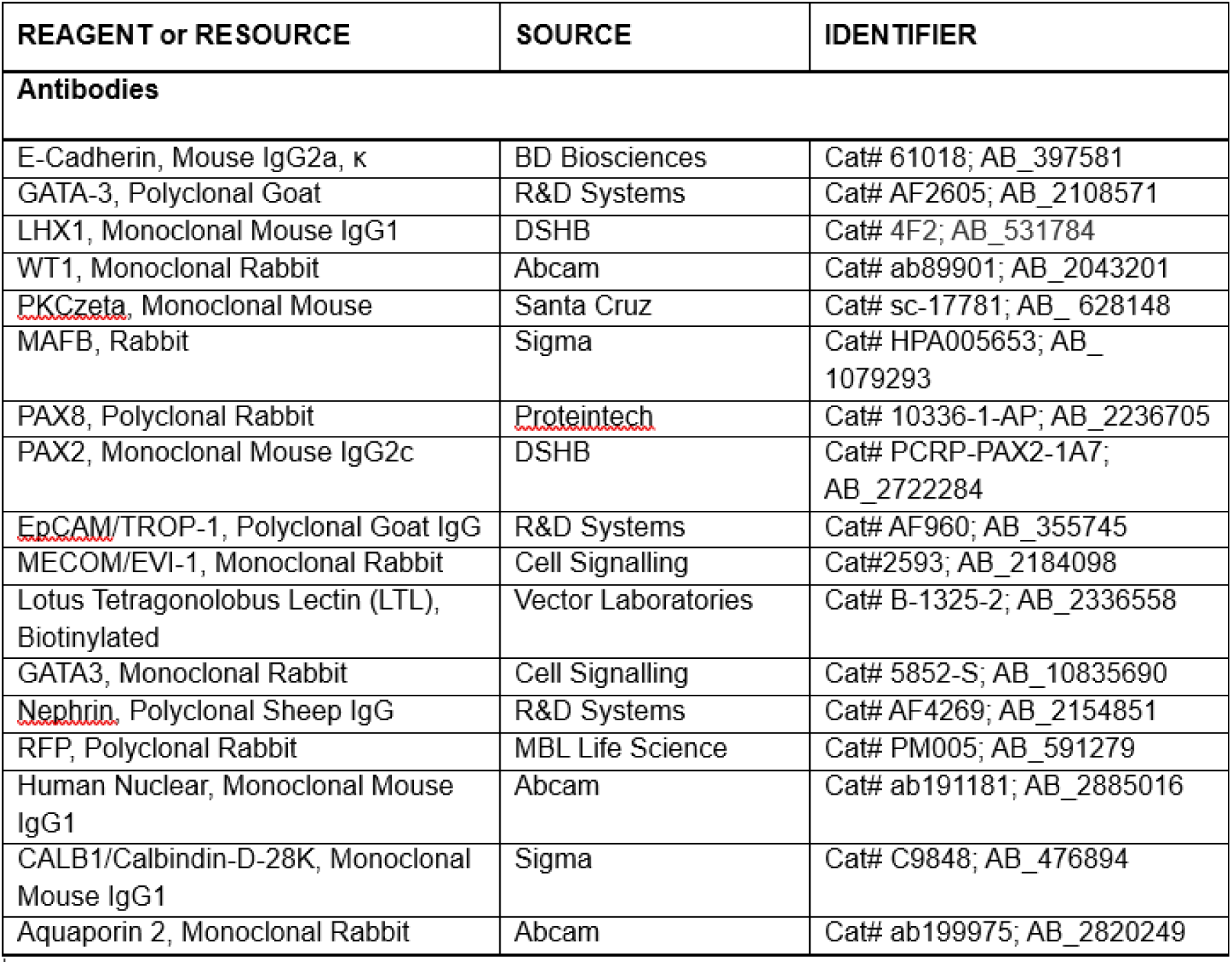
Antibodies used for immunofluorescence.

### Profiling of kidney organoids

Organoids were profiled using maximum projected images on QuPath (46) using the pixel classification tool to calculate the overall percentage of kidney organoids. The percentage of MAFB and GATA3 was calculated as the proportion of these tubules identified by immunofluorescence in the total organoid area, where DAPI equals 100%. The DAPI channel was used to segment each individual organoid using Meta’s Segment Anything Model (SAM) (43). These areas were combined with the MAFB and GATA3 masks generated above to calculated total area. Organoids were subclassified based on morphology into one of 5 classifications: proximal, proximal bias, balanced, distal bias or distal. Proximal and distal categories consisted of organoids that only contained MAFB or GATA3 respectively. Organoids with a greater than 50% difference for MAFB percentage and GATA3 percentage were classified as distal bias, while those with −50% difference were classified as proximal bias. Organoids with only one stain were classified as either proximal (MAFB/WT1 only) or distal (GATA3 only). Balanced organoids were those that fell between the −50% to 50% percentage difference. To calculate kidney size, 5 fields of view were imaged, and then quantified using the StarDist plugin (39) on ImageJ Fiji (38) to calculate area per organoid.

### Dissociation of kidney organoids

Kidney organoids were transferred into a 15ml falcon for dissociation in 4ml of 1:1 TrypLE/Accutase solution (STEMCELL Technologies) at 37°C with occasional agitation (flicking) until large clumps were no longer visible. If the organoids were not dissociated after 20 minutes, mechanical dissociation was undertaken. The TrypLE/Accutase solution was quenched using an equal amount of TeSR-E6 and cells were run through a pre-wet 40µm cell strainer to make them into single cell suspension.

### Single cell RNA-seq analysis (scRNA-seq) and dataset generation

Single cell suspensions of each condition were fixed with formaldehyde (ProSciTech C004) and additive according to the Chromium Next GEM Single Cell Fixed RNA Sample Preparation Kit (Chromium 10x Genomics). Fixation was performed at 4°C overnight, and after addition of the buffer, fixed cells were stored at −80°C until processing. The four probe barcode version of the Chromium Fixed RNA Kit, Human Transcriptome (10x Genomics) was used to barcode all 8 samples, giving 2 libraries. Post hybridisation washes were completed individually for each barcoded sample, according to the manufacturer’s protocol. Cell counting was completed using Invitrogen^TM^ ReadyCount^TM^ Green/Red Cell Viability Stain (Thermo Fisher Scientific) and the Countess 3 Automated Cell Counter (Thermo Fisher Scientific). A target of 10,000 cells per sample was combined for a total of 40,000 cells per library.

### Analysis of scRNA-seq datasets

Single cell RNA-sequencing was performed on suspension kidney organoids using the MAFB^mTagBFP2^;GATA3^mCherry^ human iPSC line. To reduce variability, libraries were created from the same monolayer thawed on the same day and analysed at two time points (D7+5 and D7+11). Organoids were made into single cell suspensions, then underwent single cell RNA-sequencing following the 10X Chromium Single Cell Flex method. At D7+5, each sample contained 10,000-13,000 individual cell transcriptomes per growth factor condition. At D7+11, each sample had 9,500-10,500 individual cell transcriptomes per growth factor. Samples were integrated using Scanorama (40) and clustered using the Leiden algorithm (42), identifying either 14 (D7+5) or 17 (D7+11) clusters. These underwent dimensionality reduction using UMAP. Clusters were classified using a combination of *DevKidCC* (25) with NPC-like reclassified as unassigned, and marker genes. These were then separated by growth factor conditions. Differential expression analysis was completed using the ScanPy’s rank genes group function method using t-tests (41).

### Kidney organoid transplantation under the murine renal capsule

All animal experiments were approved by the animal ethics committee at Murdoch Children’s Research Institute. Organoids for transplantation were generated as above, and were collected and placed in media containing 10ug/ml of the corresponding FGF (or 5ug/ml each when FGF10 and 8 were combined) to what they were grown in just prior to placement under the renal capsule. The immunodeficient strain NOD.Cg-Prkdc scid II2rg tm1WjI/SzJ/arc (NSG) mice (females only) were injected with Buprenorphine 30 minutes prior to surgery. Mice were anaesthetised using isoflurane inhalant (600ml/min, 4% induction, 2% maintenance and core body temperature was maintained at 37°C using a heat mat. The left kidney was exteriorised at the left flank, and a small incision was made into the renal capsule. Kidney organoids were placed under the capsule using a tube pipette. Pain relief was given post-surgery (Buprenorphine at 8 hours post-surgery, Meloxicam at 24- and 48-hours post-surgery). Mice were kept until 2-weeks post transplantation.

### Perfusion fixation and tissue collection

At 2 weeks post transplantation, mice were anaesthetised using isoflurane and euthanised by exsanguination. An incision was made in the abdominal wall to expose the kidneys and extended up the thoracic wall and through the diaphragm to expose the heart. A winged infusion set with a 23-gauge butterfly needle was inserted into the left ventricle and a small incision was made into the right atrium. Kidneys were cleared of blood by adding 20ml of PBS at 5ml/minute through the left ventricle and were then fixed using 20ml of 4% paraformaldehyde solution. Kidneys were subsequently immersion-fixed in 4% paraformaldehyde for a further hour.

### Vibratome Sectioning and Wholemount Immunofluorescence

Kidneys were placed in 3% low melting point analytical grade agarose (Promega, cat# V2111) and cut on a Leica vibratome (Leica VT1200) to 600um thick sections. For staining, samples were moved to a 24 well plate, and primary antibodies were diluted in 0.1% Triton-X100/PBS. Primary antibodies were detected with Alexa Fluor-conjugated fluorescent secondary antibodies (Life Technologies) at 1:500. Nuclei were detected with DAPI diluted at 1:1000 in PBS given with the secondary antibody diluent. Sections were placed in antibody diluents for 5 days, with 2 rinses and a 3-day wash between primary and secondary antibody incubations, as well as after the final secondary antibody incubation. Samples were dehydrated at 4°C in ethanol (1×50% ethanol, 2×100% ethanol) for one hour each. Samples were then cleared in ethyl cinnamate (Sigma-Aldrich) and imaged on the Andor Dragonfly 500 Spinning Disk confocal microscope (Andor Technology, Belfast, United Kingdom). Images were processed using Fiji ImageJ (38).

### Transplant analysis

Total distal tubule number and touchdown number in graft tissues were counted manually from z-stacks of 600um tissue. Total glomeruli area and total distal area were calculated from at least 3 sections within these z-stacks in a semi-automatic process involving the Napari-SAM plugin (43, 44). For total glomeruli area, this involved bounding box prompts, whereas total distal area was calculated using the Otsu’s method. Both annotations were followed by manual QC.

### Statistical Analysis

Unless otherwise stated, datasets are presented as Mean ± SEM. All datasets were analysed using GraphPad Prism version 10.0, where P<0.05 was considered significant.

## Associated Data

### Supplementary Materials

**Supplementary Figure 1:**
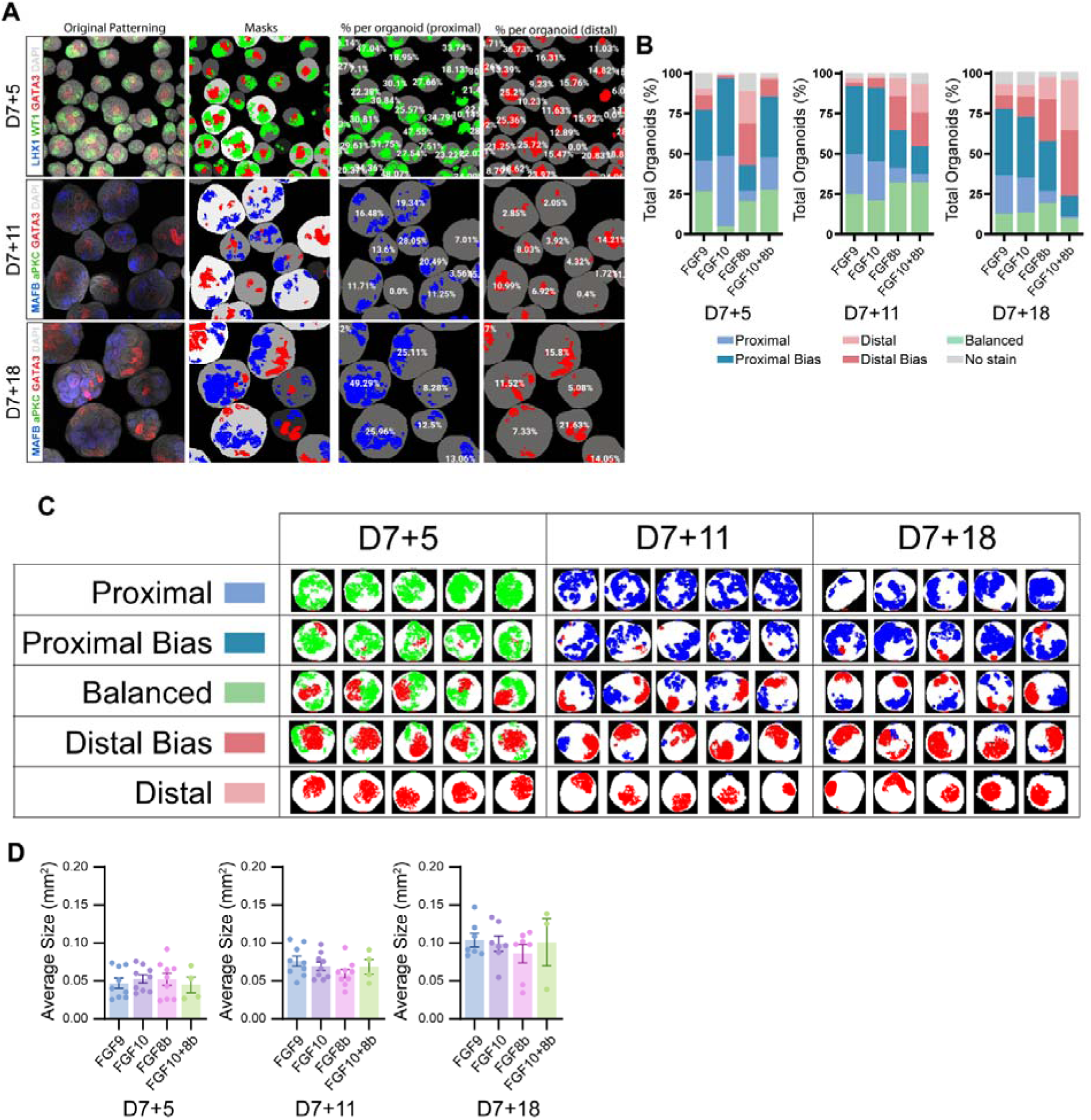
Organoid classification based on patterning differences. **A.** Organoid percentage was calculated using pixel classification tools to generate masks. These were used to generate percentage of proximal or distal proportions per organoid. **B.** Stacked bar charts showing pooled organoids subclassified based on morphology. **C.** Example organoids per subclassification used to generate pie charts and stacked bar charts, separated into subclassification per time point. **D.** Quantification of average organoid size at multiple time points, where one replicate equals the average of one differentiation. All graph data indicate mean ± SEM; *n=8-9* individual differentiations per condition per time point, except for FGF10+8b where *n=4*. A one-way ANOVA with a Dunnett’s multiple comparisons *post hoc* test was used for analysis to compare to FGF9 control.

**Supplementary Figure 2:**
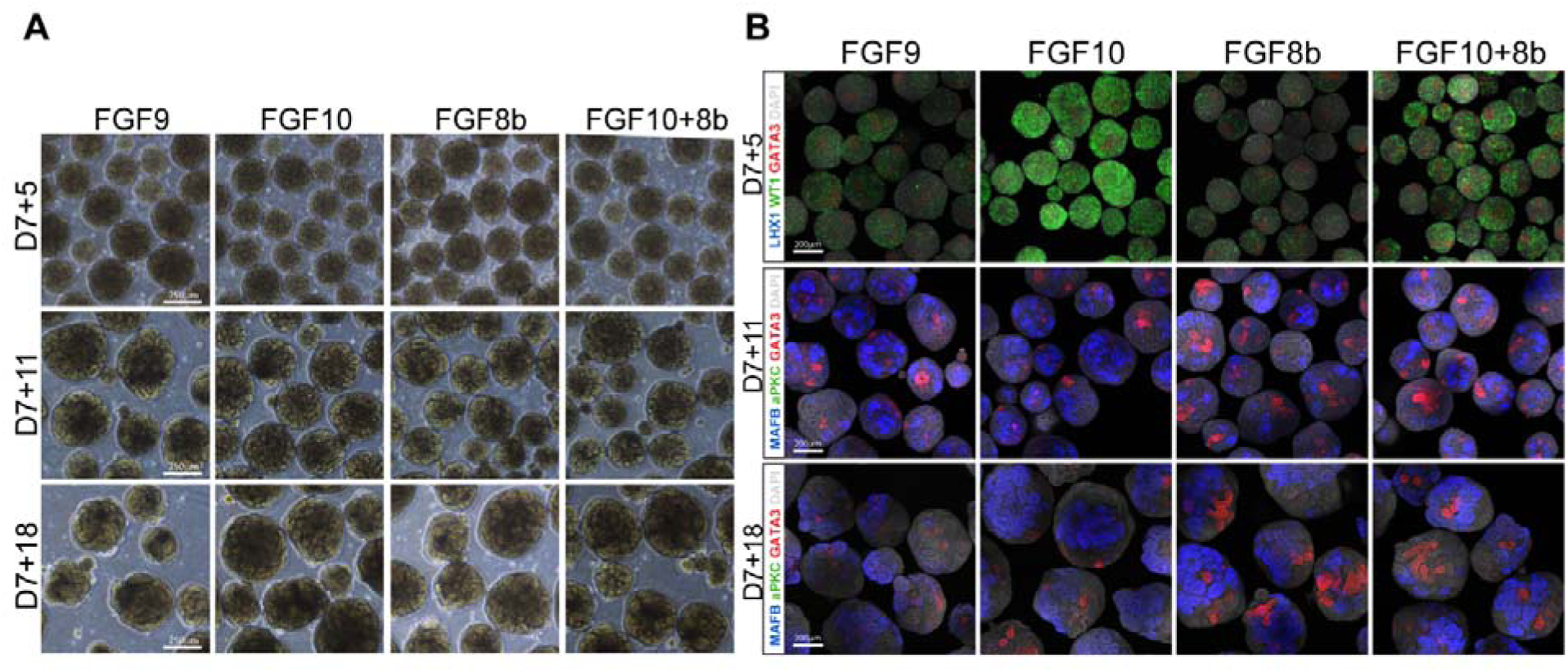
Patterning differences are seen in organoids from multiple human derived iPSC lines. **A.** Representative brightfield images of organoids following variations at multiple timepoints. This line is the CALB1^mCherry^ human iPSC cell line. **B.** Representative confocal immunofluorescence images. At D7+5, immunofluorescence depicts distal early nephrons (GATA3; red), renal vesicle (LHX1; blue) and proximal early nephron (WT1; green) and a nuclear marker (DAPI; grey). At D7+11 and D7+18, , it shows podocytes in the glomeruli (MAFB; blue), late distal tubule (GATA3; red), apical surface of the renal tubules (aPKC; green) and a nuclear marker (DAPI; grey). Scale bars = 250 µm.

**Supplementary Figure 3:**
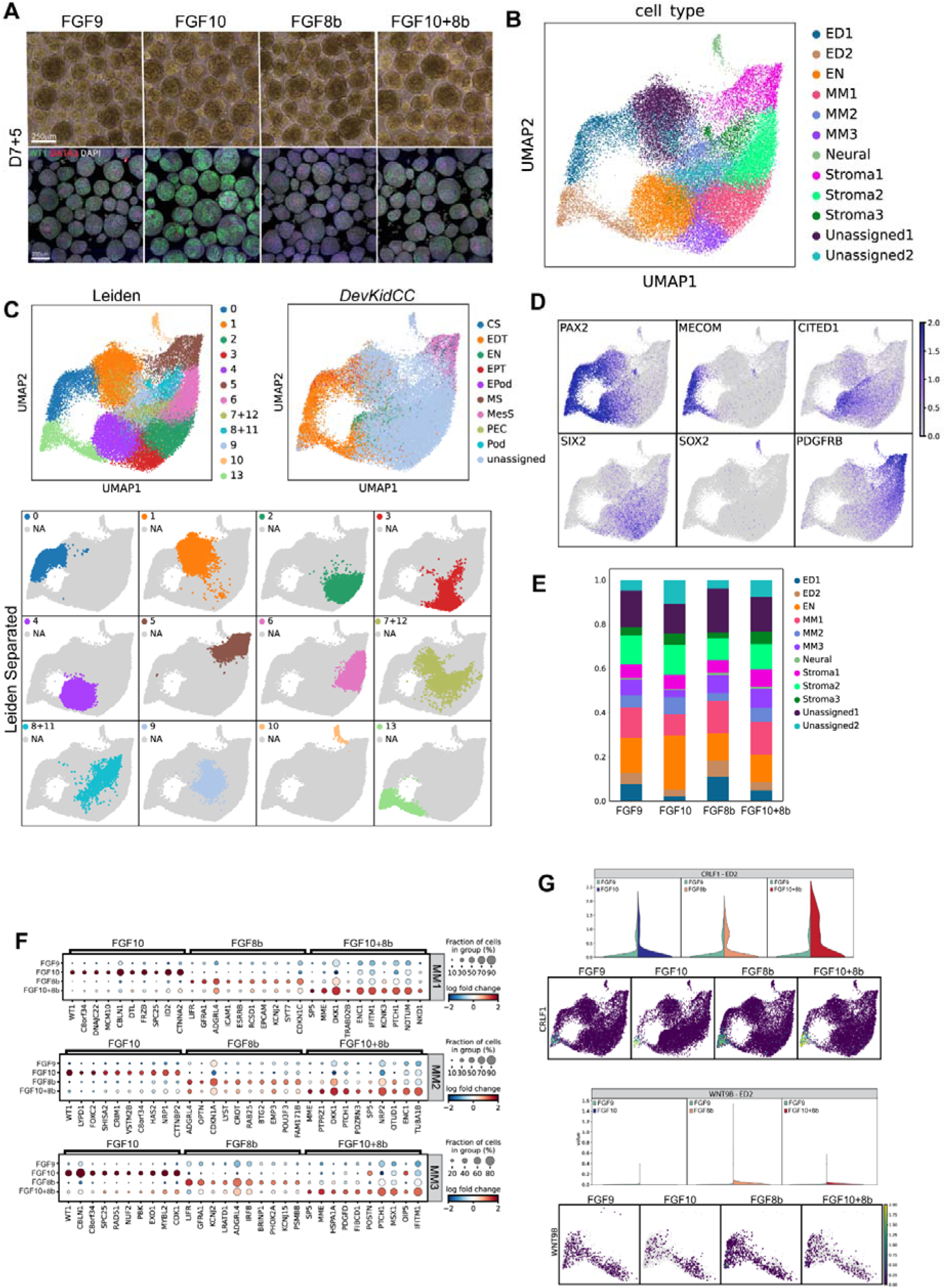
Classification of single cell analysis at D7+5. **A.** Brightfield and immunofluorescence images of organoids used for single cell analysis at D7+5, cultured in different growth factors (FGF9 control, FGF10, FGF8 and FGF10+8). Immunofluorescence depicts distal early nephrons (GATA3; red), renal vesicle (LHX1; blue) and proximal early nephron (WT1; green) and a nuclear marker (DAPI; grey). Scale bars = 200 µm. **B.** Cell type classifications, segregated by Leiden cluster and *DevKidCC* **(C)**. Unknown Leiden clusters are segregated based on marker genes **(D)**. **E.** Proportion plots showing the proportion of cell types in kidney organoids. Population abbreviations: metanephric mesenchyme (MM) separated into 3 Leiden clusters (MM1, MM2, MM3), early distal (ED) separated into two Leiden clusters (ED1, ED2), early nephron (EN). **F.** Top 10 significantly differentially expressed genes, expressed with a cutoff of >25% cells having positive expression. These are separated out into cell type clusters, showing metanephric mesenchyme Leiden clusters 1, 2 and 3. **G.** Expression of nephric duct markers CRLF1 and WNT9B in early distal cluster 2 (ED2).

**Supplementary Figure 4:**
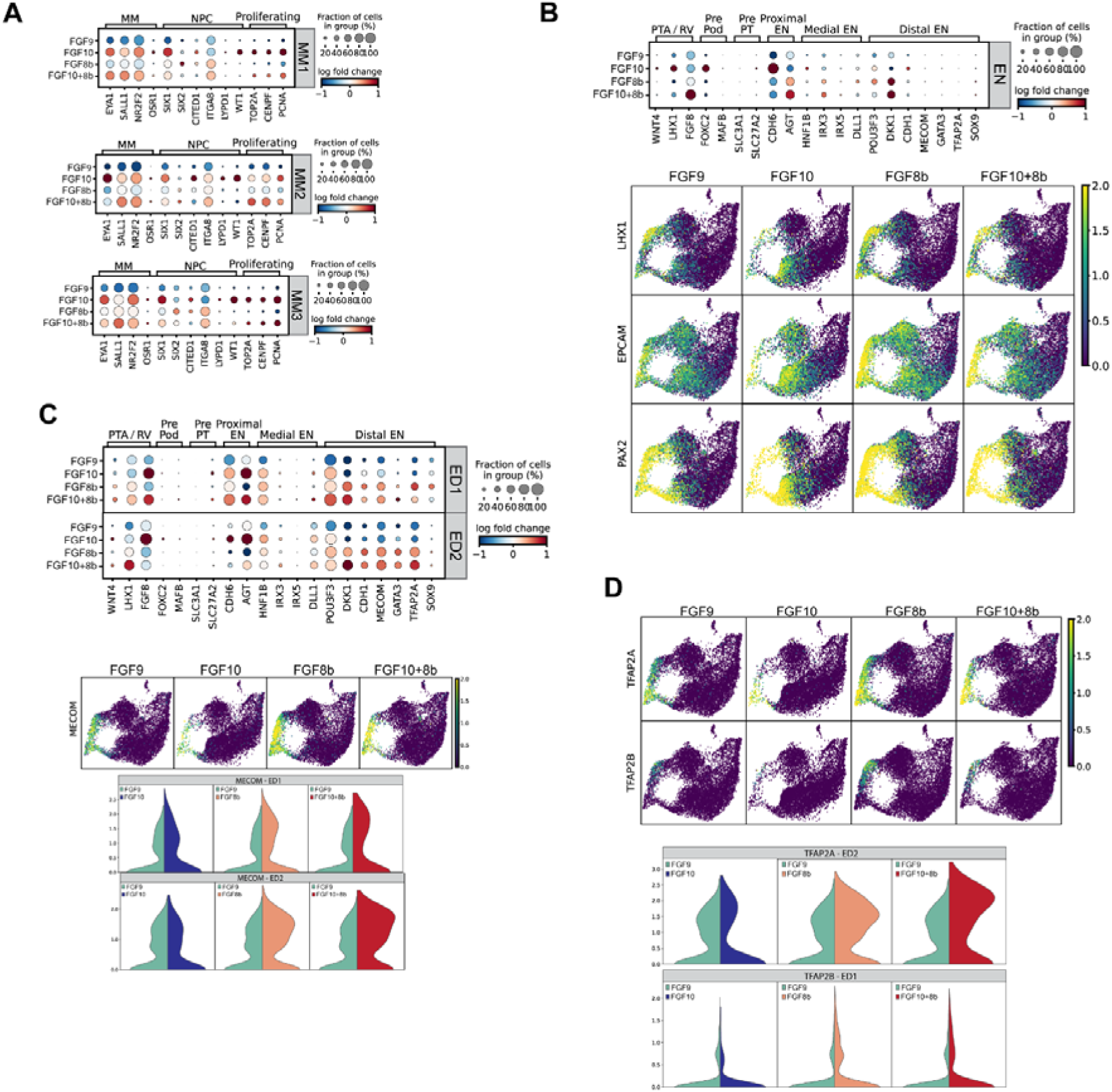
Growth factor variations retain expression of key renal segment markers early in kidney organoid development. **A.** Dot plots showing expression of marker genes, separated out via kidney segment markers by growth factor in the metanephric mesenchyme clusters (1, 2 and 3). These include markers of the metanephric mesenchyme (MM), nephron progenitor cells (NPC) and proliferating cells. **B.** Dot plots showing expression of marker genes in the early nephron (EN) cluster. Abbreviations: PTA/RV (pretubular aggregate/renal vesicle), pre-podocytes (Pre Pod), pre-proximal tubules (Pre PT), early nephron (EN). Accompanying UMAPs highlighting key genes of the PTA/RV (LHX1, EPCAM, PAX2). **C.** Dot plots showing expression of marker genes in the early distal (ED) clusters 1 and 2. Abbreviations: PTA/RV (pretubular aggregate/renal vesicle), pre-podocytes (Pre Pod), pre-proximal tubules (Pre PT), early nephron (EN). UMAPS and violin plots of both clusters, showing the early distal marker MECOM. **D.** UMAPs and violin plots of markers known to modulate nephron precursors differentiation into distal nephron (TFAP2A, TFAP2B).

**Supplementary Figure 5:**
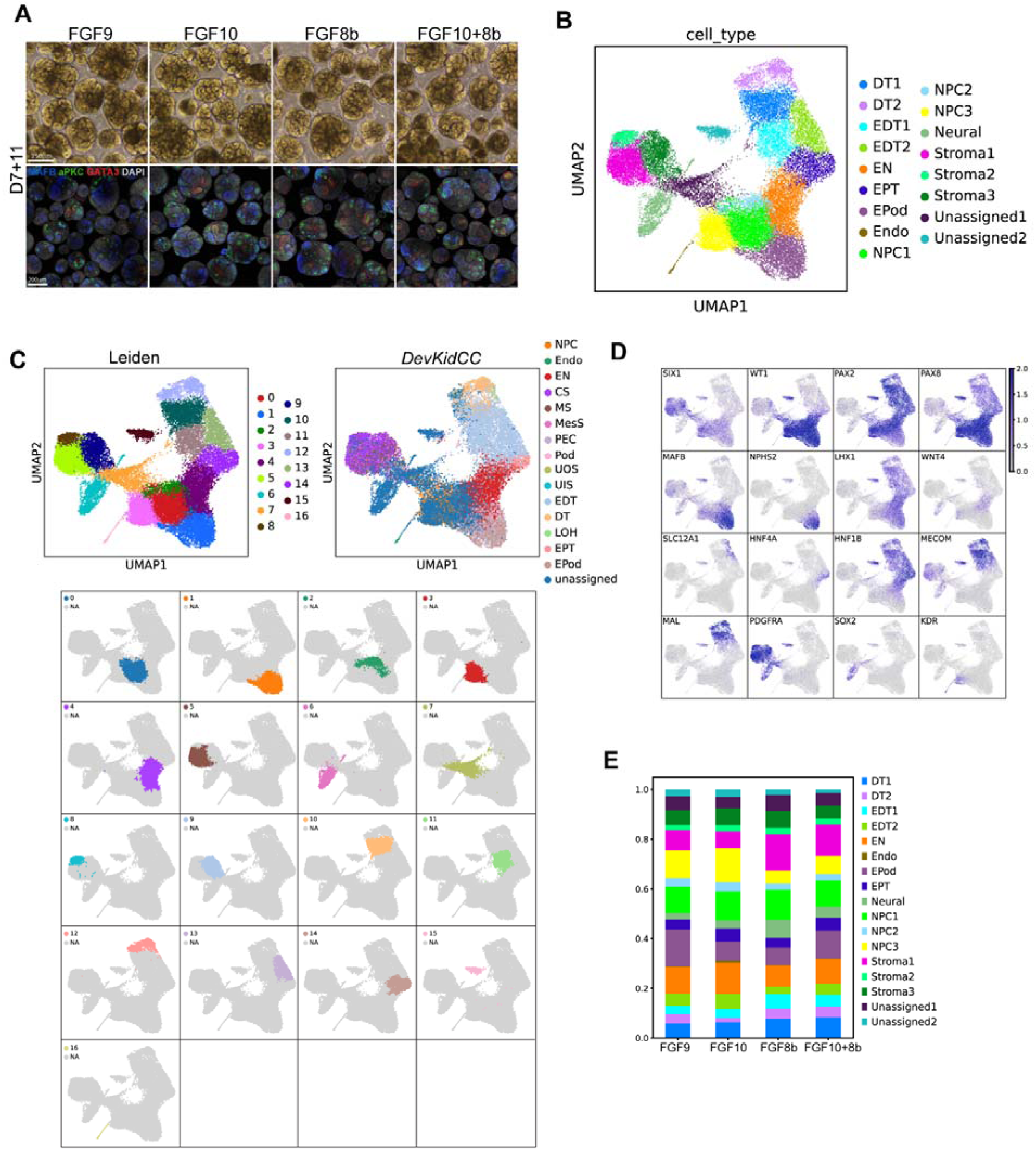
Growth factor variations retain expression of key renal segment markers late in kidney organoid development. **A.** Brightfield and immunofluorescence images of organoids used for single cell analysis at D7+11, cultured in different growth factors (FGF9 control, FGF10, FGF8 and FGF10+8). Immunofluorescence depicts podocytes in the glomeruli (MAFB; blue), late distal tubule (GATA3; red), apical surface of the renal tubules (aPKC; green) and a nuclear marker (DAPI; grey). **B.** Cell type classifications of D7+11 organoids, segregated by Leiden cluster and *DevKidCC* **(C)**. Abbreviations: nephron progenitor cell (NPC), early nephron (EN), early distal tubule (EDT), DT (distal tubule), loop of Henle (LOH), early proximal tubule (EPT), parietal epithelial cell (PEC), early podocyte (EPod), podocyte (Pod), endothelium (Endo), cortical stroma (CS), medullary stroma (MS), ureteric outer stalk (UOS), ureteric inner stalk (UIS), mesangial cells (MesS). Unknown Leiden clusters are classified based on marker genes **(D)**. **E.** Proportion plots showing the proportion of cell types in kidney organoids. Population abbreviations: nephron progenitor cell (NPC), early nephron (EN), early distal tubule (EDT), DT (distal tubule), loop of Henle (LOH), early proximal tubule (EPT), parietal epithelial cell (PEC), early podocyte (EPod), podocyte (Pod), endothelium (Endo), cortical stroma (CS), medullary stroma (MS), ureteric outer stalk (UOS), ureteric inner stalk (UIS), mesangial cells (MesS).

**Supplementary File 1:** Top differentially expressed genes for the following sample comparisons: FGF9 versus FGF10 versus FGF8b versus FGF10+8b, expressed in at least 25% of cells per cluster at D7+5. Related to Figure 3, Supplementary Figure 3, Supplementary Figure 4.

**Supplementary File 2:** Top differentially expressed genes for the following sample comparisons: FGF9 versus FGF10 versus FGF8b versus FGF10+8b at D7+11. Related to Figure 4, Supplementary Figure 5.

**Supplementary File 3:** Marker genes expressed in re-clustered Distal Tubule 1 and Distal Tubule 2 populations. Related to Figure 4.

